# Acute scalp EEG signatures of anterior thalamic DBS in epilepsy: reproducible gamma power enhancement within an aperiodic spectral-state shift

**DOI:** 10.64898/2026.10.16.759433

**Authors:** Elaheh Sabbaghi, Herkko Mattila, Eetu Siitama, Jari Hyttinen, Jukka Peltola, Narayan Puthanmadam Subramaniyam

## Abstract

**Background:** Deep brain stimulation (DBS) of the anterior nucleus of the thalamus (ANT) is an established treatment for drug-resistant epilepsy, but acute scalp-EEG signatures of stimulation remain incompletely characterized.

**Objective:** To identify acute scalp-EEG signatures of ANT-DBS and assess their cross-year reproducibility in a proof-of-concept repeated-session cohort.

**Methods:** Scalp EEG from five patients recorded one year apart (10 sessions) was analyzed during nominal cyclic DBS (1 s ON, ~5 s OFF). DBS ON was compared with the within-cycle OFF mean and consecutive post-offset OFF windows using power spectral density (PSD), spatiotemporal–frequency (STF) analysis, and spectral parameterization.

**Results:** DBS ON showed higher 30–70 Hz gamma power than OFF_mean_ in all sessions (median +0.94 dB), with significant PSD clusters overlapping gamma in 10/10 sessions. STF identified gamma increases as the most reproducible spatiotemporal effect. Exploratory frontal 34–44 Hz and central 56–70 Hz components showed cross-year support, and the broader pattern persisted when 48–52 or 45–55 Hz were excluded before clustering. Lower-frequency effects were more heterogeneous and weakened from early to late OFF windows. Aperiodic parameterization showed lower ON exponent and offset in all sessions (Δ*χ* = −0.20; Δ*b* = −0.25). After correction, theta suppression was not preserved, whereas a smaller low-gamma residual remained.

**Conclusions:** ANT-DBS was associated with consistent acute scalp-EEG state signatures comprising raw gamma enhancement, lower aperiodic exponent and offset, and a modest residual low-gamma effect. These hypothesis-generating findings show that scalp EEG can capture acute ANT-DBS state changes, while larger prospective studies are needed to establish clinical relevance.

## 1. Introduction

Drug-resistant epilepsy (DRE) remains a major clinical challenge and is defined as failure of two tolerated and appropriately chosen antiseizure medication regimens to achieve sustained seizure freedom [1]. For patients who are not candidates for resection, or who continue to have disabling seizures after surgery, neuromodulation provides an evidence-based non-resective treatment option [2, 3]. Among neuromodulation approaches, deep brain stimulation of the anterior nucleus of the thalamus (ANT-DBS) is particularly relevant for focal epilepsy because the ANT is a key node in limbic and thalamocortical circuits involved in seizure propagation and network regulation [4, 5]. The SANTE trial, long-term follow-up studies, and subsequent registry data have shown sustained clinical benefit with an acceptable safety profile [6–10]. However, although ANT-DBS is clinically established, the acute neurophysiological effects detectable noninvasively at the scalp remain incompletely defined.

Existing electrophysiological studies show that ANT-DBS modulates epileptic networks, but they leave important questions about acute scalp-EEG stimulation-state effects unresolved. Invasive recordings have linked high-frequency ANT stimulation to hippocampal desynchronization and decoupling of large-scale epileptic networks [11]. More recently, longitudinal intracranial ANT recordings over four years showed that responders developed a progressive shift toward higher-frequency activity, with increasing β_1_ and gamma and decreasing delta and theta power relative to non-responders [12]. Other studies have focused on seizure monitoring and risk tracking [13], while recent reviews have considered biomarkers of longer-term DBS response [14]. Preclinical work has similarly reported reduced theta and increased gamma power [15]. Together, these studies support the view that ANT-DBS acts through distributed epileptic and thalamocortical networks, but they do not directly define reproducible acute scalp-level spectral signatures of the DBS ON state.

Scalp EEG studies of ANT-DBS are fewer and have addressed related but distinct questions. Concurrent scalp EEG has linked effective cyclic ANT-DBS to temporal-lobe theta-band desynchronization and broader theta/alpha reductions in responders [16]. Retrospective EEG analyses have reported responder-associated decreases in beta- and gamma-band power together with reduced gamma-band phase synchronization [17]. More recently, time-locked scalp EEG analysis identified an offset-related event-related potential and corresponding limbic source activity during clinically delivered high-frequency ANT-DBS [18]. These studies indicate that scalp EEG can capture ANT-DBS-related effects. However, the immediate within-cycle spectral pattern during repeated cyclic stimulation remains insufficiently characterized. In particular, it is unclear which spectral changes are most reproducibly associated with DBS ON, which depend on the timing of the OFF reference window, and which remain stable across repeated recordings.

A further methodological issue is that raw band-power changes may not reflect isolated oscillatory modulation. Shifts in aperiodic exponent or offset can reshape apparent theta, alpha, beta, or gamma power even when narrowband activity is not selectively changed [19, 20]. This distinction is especially important in DBS–EEG because within-cycle ON–OFF contrasts occur close to stimulation transitions and may reflect both stimulation-linked changes during ON and rapid recovery after offset. Evidence that aperiodic activity is sensitive to stimulation state comes from subthalamic DBS, where the aperiodic exponent distinguished stimulation states, and from intracranial epilepsy recordings, where the exponent decreased after responsive stimulation [21, 22]. Together with the longitudinal ANT-LFP findings, these studies suggest that stimulation-related spectral states may be expressed through both higher-frequency and aperiodic changes. To our knowledge, a noninvasive scalp-EEG counterpart that captures acute ANT-DBS effects while explicitly separating periodic and aperiodic contributions has not been reported. Concurrent DBS–EEG/MEG is also vulnerable to stimulation-related artifacts, making careful interpretation essential [23, 24]. Therefore, acute ANT-DBS scalp EEG should be examined not only through raw band-power summaries, but also through analyses of post-offset timing, spatiotemporal reproducibility, and aperiodic spectral structure. To address this gap, we conducted a proof-of-concept study of scalp EEG from five ANT-DBS patients recorded twice, one year apart, allowing us to examine within-patient cross-year consistency. We asked which scalp-EEG changes were reproducibly associated with DBS ON, how these changes evolved across the first five seconds after stimulation offset, and how their interpretation changed after separating fitted aperiodic and residual band-limited spectral components. We combined frequency-resolved PSD analysis, sensor-level spatiotemporal–frequency analysis, and spectral parameterization to characterize acute stimulation-related spectral dynamics. Our aim was to identify and characterize acute scalp-EEG signatures of ANT-DBS, assess their cross-year reproducibility, and provide a framework for future source-level, connectivity-based, and prospective validation studies.

## 2. Materials and methods

### 2.1. Participants, ethics, and design

We analyzed scalp EEG from five adults with drugresistant focal epilepsy treated with deep brain stimulation of the anterior nucleus of the thalamus (ANT-DBS) at Tampere University Hospital. The study was approved by the Ethics Committee of the Pirkanmaa Hospital District (decision R17132; approved 19 September 2017). All participants provided written informed consent, identifiers were removed before analysis, and the study followed the Declaration of Helsinki [25]. Each patient was recorded in 2018 and 2019, yielding 10 patient-year sessions for within-patient cross-year assessment. The cohort was heterogeneous in age, epilepsy duration, etiology, and epileptogenic-zone distribution. The study characterized acute stimulation-state EEG effects rather than therapeutic efficacy. There was no randomization, blinding, or separate sham condition; within-cycle DBS OFF periods served as the physiological comparator. Clinical characteristics are summarized in Table 1, with stimulation-programming details in the Supplementary Material.

**Table 1.** Clinical characteristics of the study cohort. Age and epilepsy duration are reported at each patient’s 2018 DBS–EEG recording. Epilepsy duration was calculated from age at the 2018 recording and age at epilepsy onset. The predominant seizure type was FIAS in all patients. M, male; F, female; CD, cortical dysplasia; FIAS, focal impaired awareness seizure.

| Patient | Age (y) | Sex | Epilepsy duration (y) | Onset age (y) | Etiology | Epileptic zone |
| --- | --- | --- | --- | --- | --- | --- |
| P01 | 53 | M | 52 | 1 | Ischemic lesion (right parietal) | Multilobar (frontoparietal) |
| P02 | 60 | M | 46 | 14 | Unknown | Multifocal |
| P03 | 33 | F | 10 | 23 | Viral encephalitis | Multifocal |
| P04 | 24 | M | 6 | 18 | CD; bilateral subependymal periventricular heterotopy | Multifocal |
| P05 | 62 | M | 44 | 18 | CD | Unifocal multilobar (left frontotemporal) |

Recordings comprised repeated nominal 1 s ON / ~5 s OFF cycles. Each ON window was compared with five consecutive 1 s OFF windows (OFF_1_–OFF_5_) to characterize the DBS ON state and early post-offset dynamics (Fig. 1).

**Figure 1:**
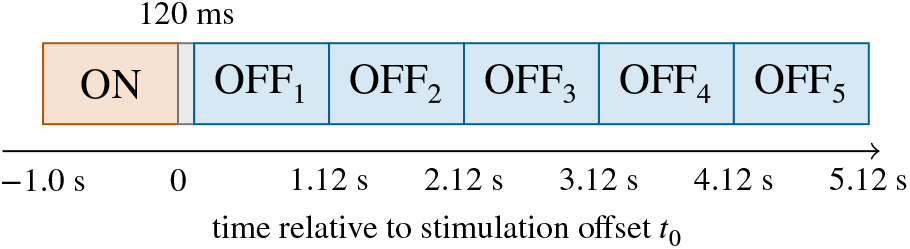
Within-cycle analysis design. The schematic shows the analyzed windows relative to stimulation offset (*t*_0_ = 0). The ON window spanned −0.98 to 0.02 s. After a 120 ms guard interval, OFF_1_–OFF_5_ covered 0.12–5.12 s. The next stimulation burst began after OFF_5_ had ended. OFF_mean_ denotes the mean of OFF_1_–OFF_5_ when at least 3 windows were valid.

### 2.2. EEG acquisition and stimulation protocol

EEG acquisition followed Mattila et al. [18] using a 64-channel 10–10 cap. All patients had Medtronic Activa PC pulse generators and received voltage-controlled bilateral monopolar ANT stimulation at 5.0 V per side, 140 Hz, and 90 µs during nominal 1 s ON / 5 s OFF cycles for approximately 10 min. Settings were unchanged across years. This acute time-locked paradigm preserved the 1:5 duty ratio of conventional 1-min ON / 5-min OFF cycling while providing repeated transitions during a short recording; it was not intended to reproduce chronic therapeutic cycling. Leads and contacts were reconstructed in MNI space and visualized relative to the bilateral ANT labels from the DISTAL atlas using Lead-DBS v3 (Supplementary Fig. S1; Supplementary Methods). This visualization was descriptive, and no quantitative spatial analysis was performed.

### 2.3. Preprocessing and offset-locked windowing

Continuous EEG was visually screened, and bad channels were marked before 64-component InfoMax ICA was estimated in Brainstorm after resampling to 2 kHz [18, 26]. Components reflecting ocular, small-amplitude muscle, or stimulation-related activity were removed by visual inspection of their time courses, scalp topographies, and the EEG distortion produced by their removal. The cleaned continuous data were retained at 2 kHz for cycle extraction.

The source recordings had different prior filtering histories: the 2018 data had been filtered over 0.5–100 Hz, whereas the 2019 data had been filtered over 0.5–80 Hz with an additional 50-Hz notch. We therefore applied the same MNE `spectrum_fit` 50-Hz notch to all sessions, rereferenced each 1-s window to the common average of channels valid for that cycle, restricted analyses to 4–70 Hz, and compared ON and OFF windows from the same cleaned recording. Each 7 s epoch was time-locked to stimulation offset (*t*_0_ = 0). The ON window spanned −0.98 to 0.02 s, followed after a 120 ms guard by five consecutive 1 s OFF windows from 0.12 to 5.12 s. The next burst began at approximately 5.13 s. OFF_mean_ required at least three valid windows; further details are provided in the Supplementary Methods.

### 2.4. Spectral features

Power spectral density (PSD) was estimated for each channel and window using multitaper DPSS methods over 4–70 Hz and expressed in dB [27]. Delta activity (< 4 Hz) was excluded because 1 s windows provide limited reliability at very low frequencies. Band summaries comprised theta (4–8 Hz), alpha (8–12 Hz), beta (12–30 Hz), and gamma (30–70 Hz) and were interpreted as broad spectral ranges because of the short windows and multitaper smoothing.

Spatiotemporal–frequency (STF) representations were computed with complex Morlet wavelets from 4 to 70 Hz in 1 Hz steps [28]. The primary STF analysis retained the complete notch-cleaned frequency range. To assess dependence on the line-frequency neighborhood, cluster inference was repeated after excluding 48–52 Hz and, more conservatively, 45–55 Hz before cluster formation. Line-noise-robust gamma summaries used 30–44 and 56–70 Hz. Sensors were grouped into five macro-regions for visualization and descriptive ROI summaries, but these regions did not define the cluster-permutation tests.

Spectra were parameterized with `specparam` using a fixed aperiodic model, log_10_ *P* (*f*) = *b* − *χ* log_10_(*f*), fit over 4– 70 Hz after interpolation across 49–51 Hz in log_10_-power space [19]. Models allowed up to six peaks (width 2–12 Hz; minimum height 0.05; threshold 2.0), and channel–cycle estimates required an aperiodic-line *R*^2^ ≥ 0.20. Residual spectra were obtained after removing the fitted aperiodic component; further quality-control details are provided in the Supplementary Methods.

### 2.5. Statistical analysis

Stimulation effects were estimated within each session using cycles as paired observations. PSD and aperiodic analyses used ON–OFF_mean_ as the primary contrast, whereas STF tested ON–OFF_1_ through ON–OFF_5_ separately. Two-sided sign-flip cluster tests used 5000 permutations [29–31]. Cluster-forming nodes required an uncorrected *p* < 0.05 and at least 10 valid paired cycles for PSD and aperiodic-derived metrics or 8 for STF; sensor adjacency was defined by a 5-cm Euclidean distance. A shared maximum-cluster null controlled family-wise error across the five OFF references. Across sessions, sign replication, prevalence, and cross-year consistency were summarized descriptively; the 10 patient-year sessions were not treated as independent population-level observations. Full procedures are provided in the Supplementary Methods.

### 2.6. Software

Analyses were performed in Python 3.13.9 using MNE-Python 1.11.0 [32] and `specparam` 2.0.0rc7 [19].

## 3. Results

### 3.1. Dataset yield and quality control

We analyzed five ANT-DBS patients in two yearly sessions (2018/2019), yielding 10 patient-year sessions and 973 retained cycles (467 in 2018; 506 in 2019; 90–106 per session). Each cycle provided one ON and five OFF windows. PSD and aperiodic analyses used Δ = ON − OFF_mean_, whereas STF used separate ON–OFF_*k*_ contrasts. Session-wise QC yield is provided in the Supplementary Material. We first present raw spectral changes, then their spatiotemporal and cross-year reproducibility, and finally the fitted aperiodic and residual components.

### 3.2. Spectral power modulation during DBS (PSD, 4–70 Hz)

We first tested whether DBS ON was associated with consistent raw spectral-power changes relative to the withincycle OFF state. PSD analysis showed reduced low-frequency power and increased gamma-range power during DBS ON (Fig. 2). Across 4–70 Hz, the median ON–OFF_mean_ contrast crossed from negative to positive at approximately 24 Hz (Fig. 2A). The strongest median decrease occurred near 6 Hz (median Δ = −0.90 dB; negative in 8/10 sessions), whereas the strongest median increase occurred near 58 Hz (median Δ = +1.33 dB; positive in 10/10 sessions). Within-session channel-by-frequency cluster-permutation tests identified significant clusters overlapping gamma and beta frequencies in 10/10 sessions and theta and alpha frequencies in 8/10 sessions (Table 2).

**Table 2.**
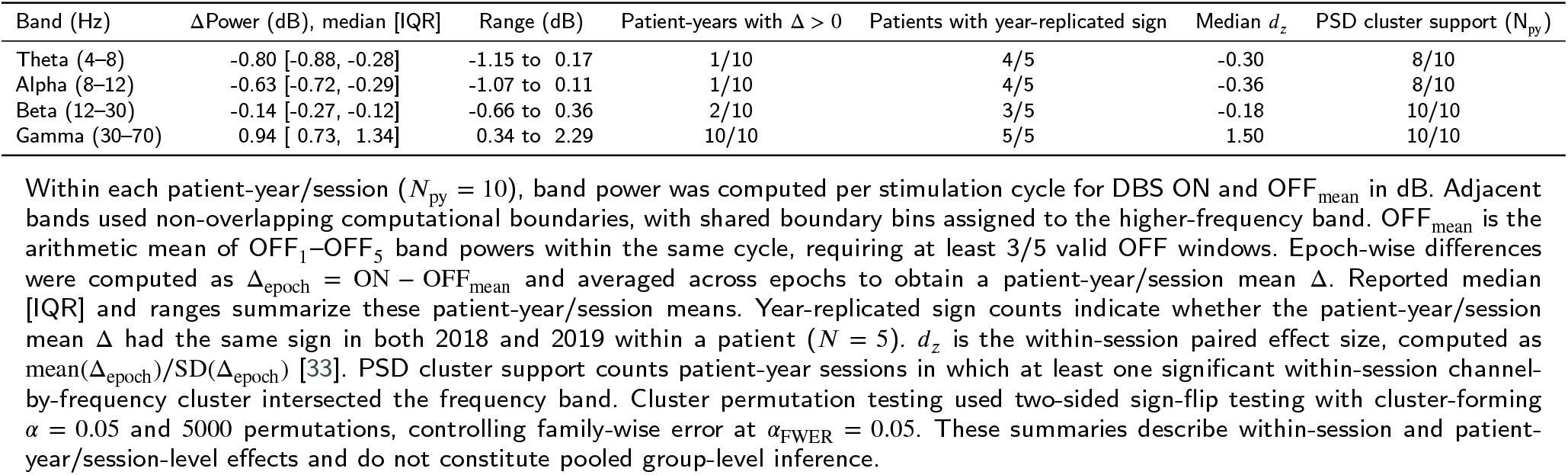
Summary of band-limited power modulation during ANT-DBS.

**Figure 2:**
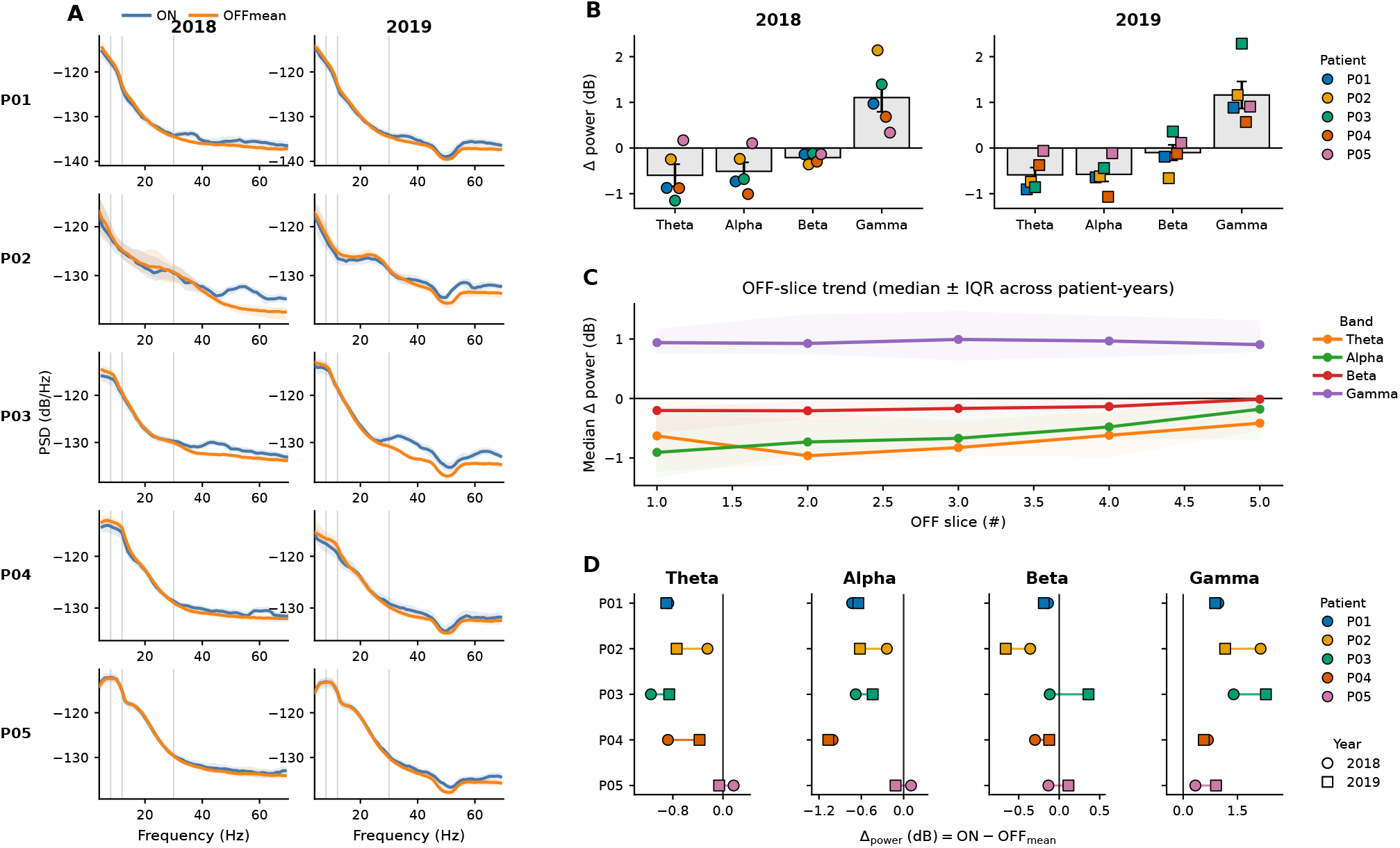
Spectral summary of stimulation-associated modulation (patient-year/session unit; *N*_py_ = 10, *N* = 5 patients). (A) PSD overlays for each annual session: DBS ON versus OFF_mean_. Curves show the median channel-mean PSD across retained cycles within each annual session; shaded bands indicate the interquartile range across cycles. (B) Band-limited changes Δ (ON−OFF_mean_) in 2018 and 2019. Points are patient-year/session mean Δ values; bars show the descriptive mean ± SEM across patients within each year. (C) OFF-window trend across OFF_1_–OFF_5_. Lines show the median Δ_*k*_ (ON−OFF_*k*_), *k* = 1, …, 5, across the 10 patient-year sessions; shaded bands show the interquartile range. (D) Patient-wise cross-year comparisons. For each band, dumbbells show each patient’s mean Δ in 2018 and 2019. OFF_mean_ denotes the within-cycle mean of OFF_1_–OFF_5_. In panels B and D, colors identify patients consistently across figures, while marker shape identifies recording year.

At the band level, gamma was the most reproducible raw PSD effect (Table 2; Fig. 2B). It increased in all sessions (median Δ = +0.94 dB [IQR 0.73, 1.34]; year-replicated sign in 5/5 patients), with a median within-session paired effect size of *d*_*z*_ = 1.50, and remained positive after 5-cycle block averaging (median +0.96 dB [IQR 0.73, 1.33]). Theta, alpha, and beta generally showed negative raw ON–OFF differences, but their reproducibility was weaker. Beta had within-session cluster support in 10/10 sessions, whereas theta and alpha were more heterogeneous across patients and recording years. However, beta effects were smaller in magnitude and showed weaker cross-year sign replication than gamma; therefore, gamma was emphasized as the more reproducible raw spectral feature.

We then examined whether ON–OFF effects depended on the timing of the OFF reference window. Low-frequency ON–OFF_*k*_ differences were largest for early OFF windows and weakened toward OFF_5_ (Fig. 2C), suggesting short-timescale post-offset dynamics rather than a fixed ON– OFF difference. This trend was clearest in alpha, followed by beta and theta, whereas gamma showed minimal OFF-window progression. Cross-year comparisons showed preserved gamma enhancement in every patient, while lower-frequency decreases were less consistent (Fig. 2D).

### 3.3. Spatiotemporal–frequency analysis and cross-year reproducibility

Having established the raw PSD pattern, we next examined how the ON–OFF differences unfolded across time, frequency, and scalp location, and whether they were reproduced one year later. The patient-year STF maps showed a broadly similar organization across sessions: lower-frequency activity tended to decrease during DBS ON, whereas gamma-range activity increased (Fig. 3A). This pattern became clearer at the cohort level, where the median and sign-consistency maps identified gamma enhancement as the most consistent STF feature (Fig. 3B). Cross-year scalp maps showed positive support both below and above the line-noise neighborhood, at 30–44 and 56–70 Hz, while negative support was concentrated mainly at lower frequencies (Fig. 3C). When these two line-noise-safe gamma ranges were combined, the mean effect was positive in all 10 sessions and retained the same direction across years in all five patients. Theta effects were less consistent: they were negative in 9/10 sessions and reproduced across years in 4/5 patients (Fig. 3D).

**Figure 3:**
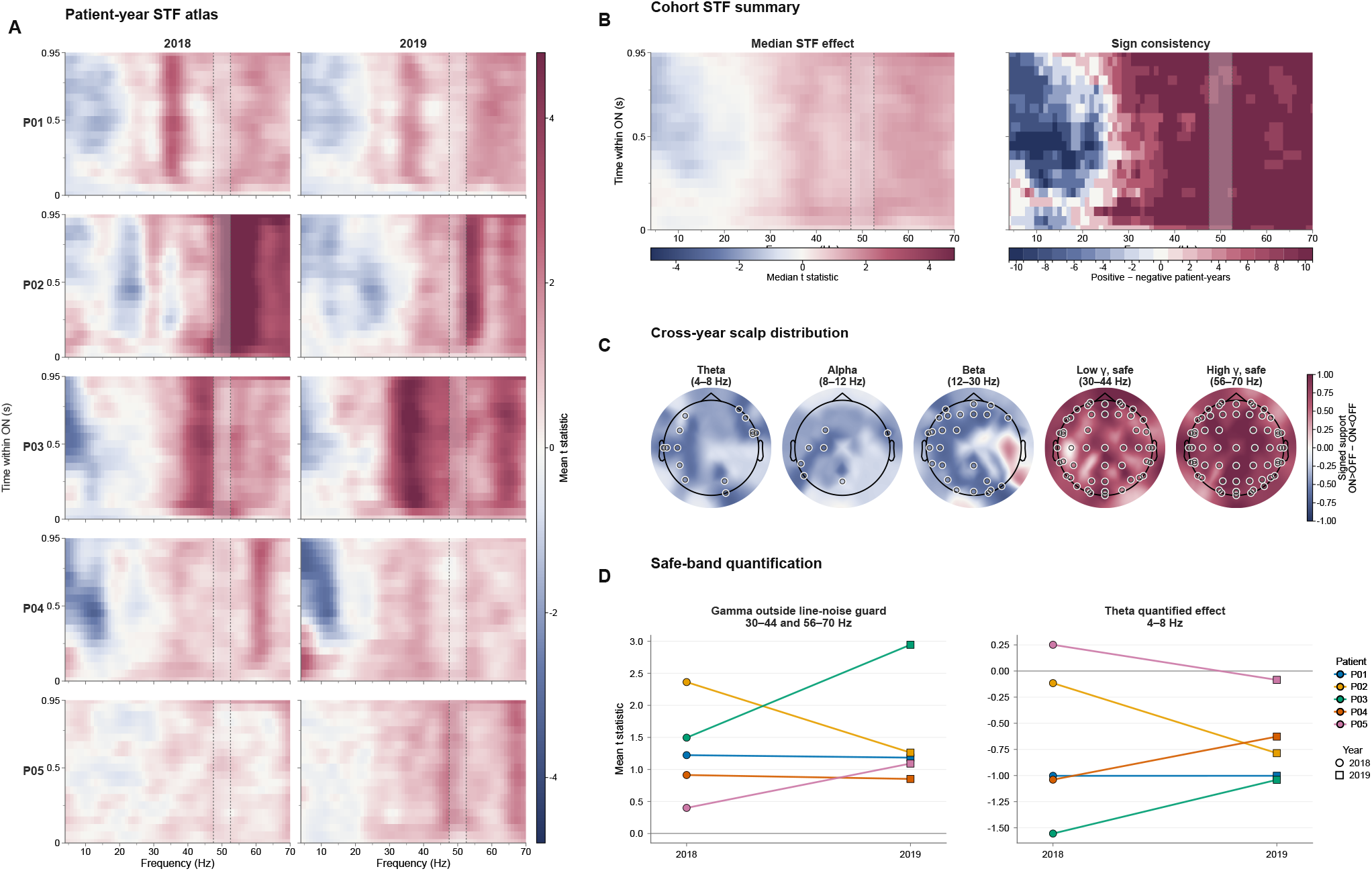
Patient-year and cohort-level STF overview of DBS ON–OFF effects. (A) Patient-year mean within-session STF *t*-statistic maps, averaged across sensors and ON–OFF_1_ through ON–OFF_5_. Warm colors indicate ON>OFF and cool colors ON<OFF. (B) Cohort median effect and sign consistency across the 10 patient-year sessions. Sign consistency is the number of positive minus negative sessions at each frequency–time bin. (C) Scalp distribution of cross-year replicated signed support. Values are the fraction of patients with replicated ON>OFF support minus the fraction with replicated ON<OFF support; open circles mark sensors with |support|≥ 3/5. (D) Descriptive within-session summaries for gamma outside the conservative guard (30–44 and 56–70 Hz) and theta (4–8 Hz). Lines connect recording years within patients. All panels use the primary notch-only analysis; the lightly shaded 48–52 Hz interval marks the conventional line-noise neighborhood for display only. Atlas and cohort maps were smoothed for display only.

OFF-reference analysis showed distinct temporal profiles (Fig. 4A,B). Gamma ON>OFF support outside 45– 55 Hz remained high across OFF_1_–OFF_5_ (95.0–97.8%), whereas 4–15 Hz ON<OFF support fell from 67.3% to 27.5%. Thus, lower-frequency effects depended more on the post-offset reference, while gamma remained comparatively stable.

**Figure 4:**
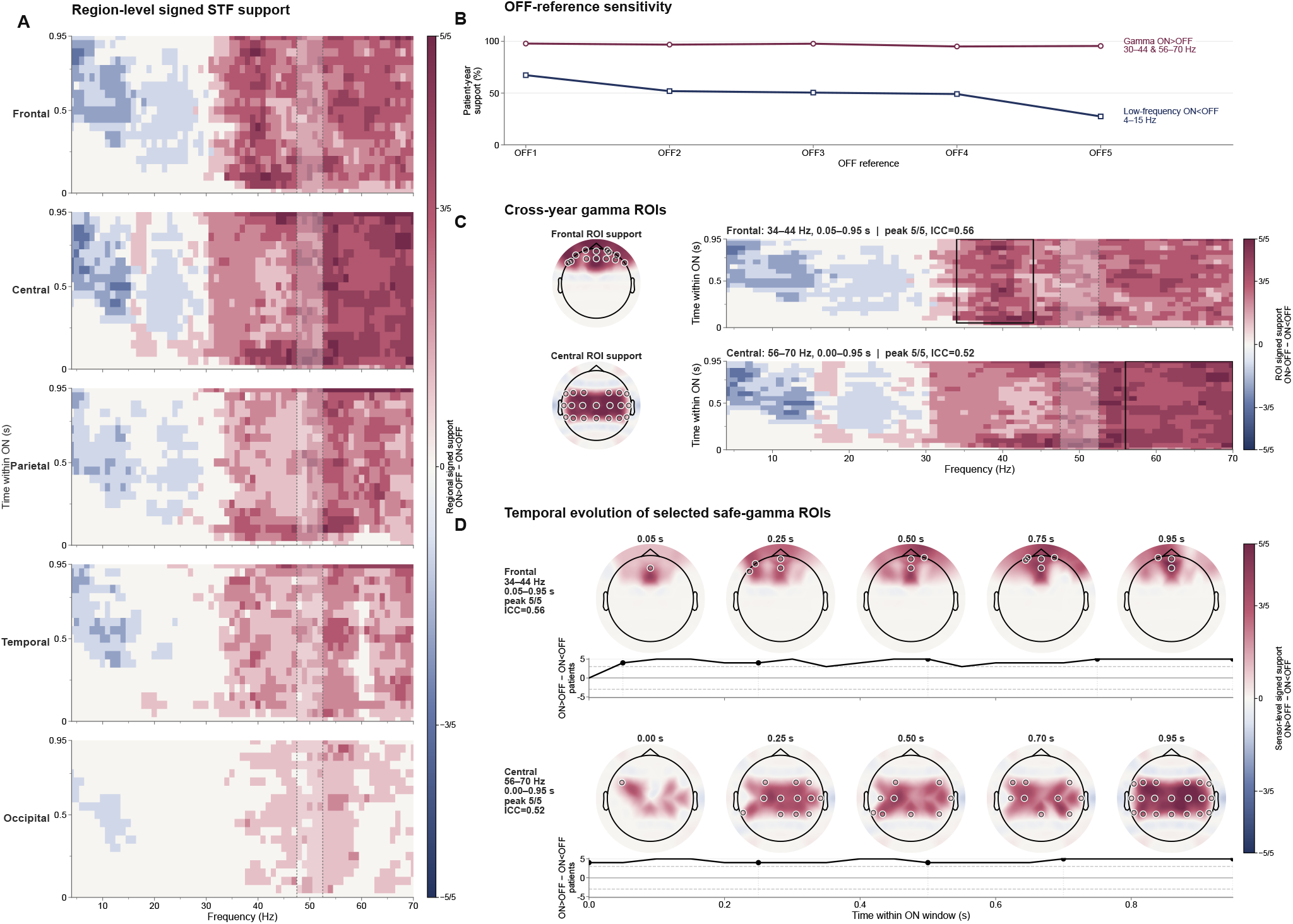
Region-level STF support, OFF-reference sensitivity, and exploratory gamma ROIs. (A) Region-level cross-year replicated signed support from the primary notch-only analysis. Warm colors indicate ON>OFF and cool colors ON<OFF. (B) OFF-reference sensitivity for gamma ON>OFF outside 45–55 Hz and low-frequency ON<OFF over 4–15 Hz. (C) Selected line-noise-safe gamma ROIs. Scalp maps show support collapsed across each fixed ROI, and rectangles show its frequency–time boundaries. ICC(3,1) values are descriptive cross-year consistency estimates calculated over the same fixed ROI in all five patients. (D) Temporal evolution of the selected ROIs. Open circles mark sensors meeting |support| ≥ 3/5. The lightly shaded 48–52 Hz interval is a display marker only. Guarded analyses were used as sensitivity tests and are shown in Supplementary Fig. S2.

We next summarized the most reproducible gamma patterns as exploratory ROIs. These were derived from the primary notch-only analysis but constrained to frequencies outside the conservative line-noise neighborhood. The selected components were a frontal 34–44 Hz pattern extending from 0.05 to 0.95 s and a central 56–70 Hz pattern extending across the full 0.00–0.95 s window. Sign-level cross-year support reached 5/5 patients for both ROIs, whereas descriptive ICC(3,1) values were moderate (0.56 and 0.52, respectively; Table 3; Fig. 4C,D). Because these ROIs were selected from the same data used to assess their consistency, they should be interpreted as descriptive summaries rather than independent validation results.

**Table 3.**
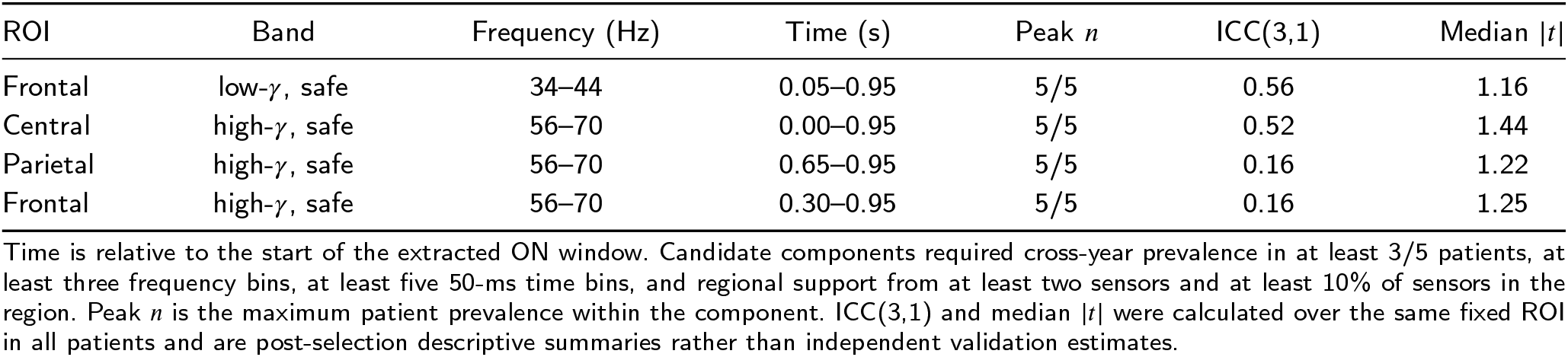
Eligible cross-year replicated gamma ROIs from the primary notch-only STF analysis. Candidates were constrained to 30–44 or 56–70 Hz. The frontal low-gamma and central high-gamma components were selected for Fig. 4 as the highest-ranked positive candidate in each interval.

Finally, the line-noise sensitivity analyses closely reproduced the primary pattern. Excluding 48–52 or 45–55 Hz retained 99.5% and 98.8% of positive nodes in the common line-noise-safe gamma domain. Gamma support remained in 50/50 and 49/50 patient-year/OFF comparisons, respectively (Supplementary Fig. S2). Together, these findings support a broad gamma-range enhancement that extends below and above the 50-Hz neighborhood, rather than a narrow effect confined to frequencies immediately adjacent to line noise.

### 3.4. Aperiodic changes and corrected spectra

Because raw PSD changes can reflect both band-limited and aperiodic components, we next separated fitted aperiodic changes from residual spectral effects. Both aperiodic parameters decreased in all 10 sessions (Fig. 5B,C; Table 4). The exponent decreased (median Δ*X* = −0.200 [IQR −0.272, −0.131]), indicating a flatter fitted spectrum, and the offset decreased (median Δ*b* = −0.254 [IQR −0.301, −0.131] log_10_-power), indicating a lower fitted spectral intercept.

**Table 4.** Aperiodic parameters and raw versus aperiodic-adjusted band-power modulation during ANT-DBS. Patient-year/session summaries (*N*_py_ = 10) describe DBS ON relative to OFF_mean_, reported as Δ = ON − OFF_mean_.

| Metric | $\Delta$ median [IQR] | $\Delta > 0$ | Replicated sign | Median $d_z$ |
| --- | --- | --- | --- | --- |
| *Aperiodic parameters* |  |  |  |  |
| Aperiodic exponent $\chi$ | -0.200 [-0.272, -0.131] | 0/10 | 5/5 | -0.85 |
| Aperiodic offset | -0.254 [-0.301, -0.131] (-2.54 [-3.01, -1.31] dB) | 0/10 | 5/5 | -0.64 |
| *Raw and aperiodic-adjusted band power* |  |  |  |  |
| Raw theta power (4–8 Hz) | -0.83 [-0.89, -0.30] dB | 1/10 | 4/5 | -0.31 |
| Aperiodic-adjusted theta (4–8 Hz) | 0.17 [0.12, 0.26] dB | 9/10 | 4/5 | 0.28 |
| Raw low-gamma power (30–55 Hz) | 0.91 [0.70, 1.17] dB | 10/10 | 5/5 | 1.16 |
| Aperiodic-adjusted low-gamma (30–55 Hz) | 0.12 [0.01, 0.21] dB | 7/10 | 4/5 | 0.52 |
| Raw high-gamma power (55–70 Hz) | 1.07 [0.94, 1.44] dB | 10/10 | 5/5 | 1.65 |
| Aperiodic-adjusted high-gamma (55–70 Hz) | -0.05 [-0.16, 0.15] dB | 4/10 | 3/5 | -0.06 |

**Figure 5:**
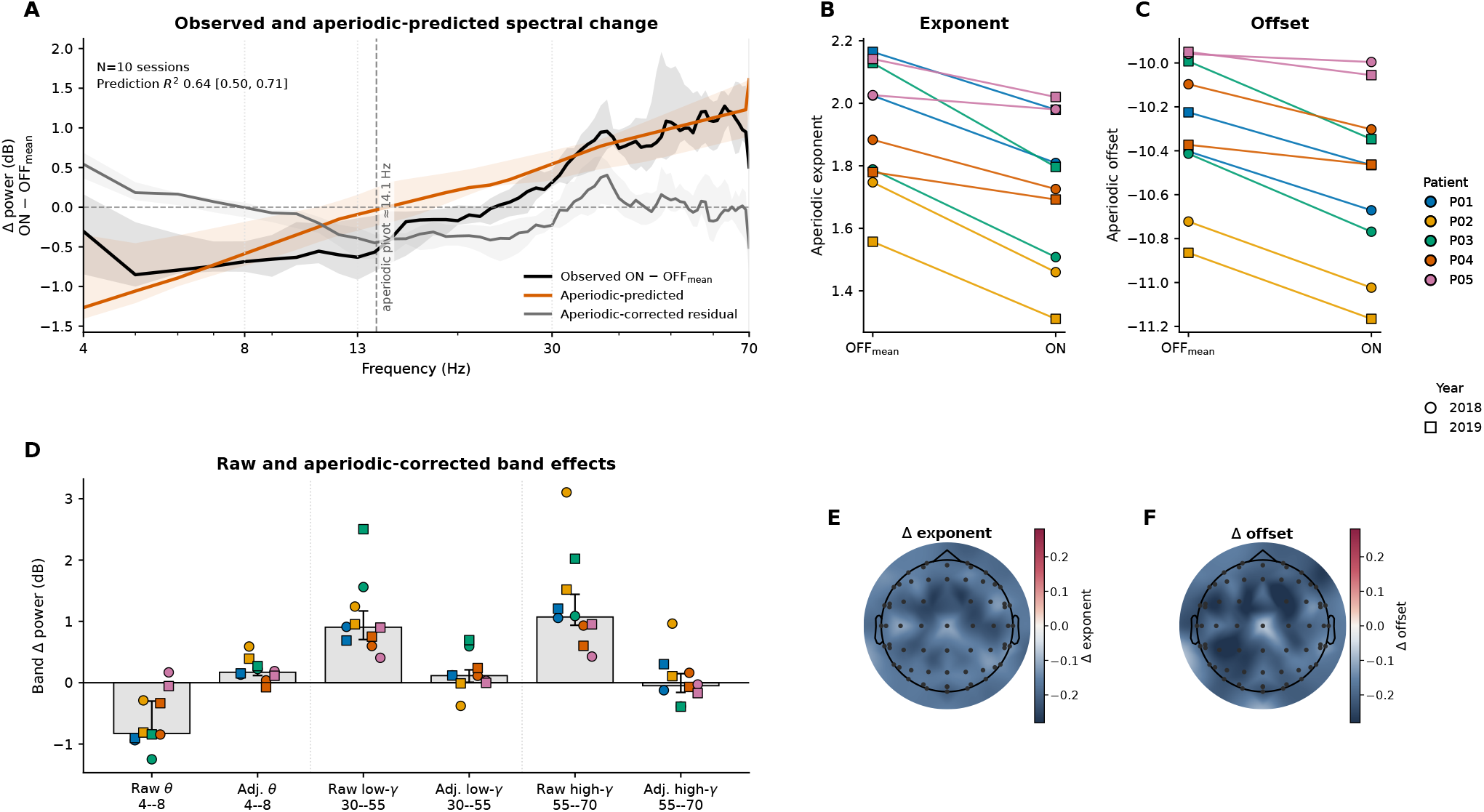
Aperiodic and residual spectral components of the DBS ON effect (*N*_py_ = 10; patient-year/session unit). (A) Observed ON–OFF_mean_ spectral change expressed in decibels as 10 ×Δ log_10_ PSD(*f*), the change predicted from the fitted aperiodic exponent and offset, and the residual after subtraction of the aperiodic prediction. Observed and predicted contrasts used the same fit-qualified channel–cycle observations. Curves show the patient-year median and interquartile range on a common 1-Hz frequency grid. The 49–51 Hz region was excluded when calculating prediction *R*^2^. The dashed vertical line marks the median aperiodic pivot frequency, at which the predicted change crosses zero. Prediction *R*^2^ is distinct from the fit-quality *R*^2^ used during spectral parameterization. (B,C) Paired OFF_mean_ and ON values for aperiodic exponent (B) and offset (C), with lines connecting conditions within each session. Offset is shown in log_10_-power units. (D) Raw and aperiodic-adjusted theta, low-gamma, and high-gamma changes. Bars show the median, error bars show the interquartile range, and markers show individual patient-year/session values. (E,F) Median sensor-level ON–OFF_mean_ changes in exponent (E) and offset (F) across patient-year sessions. Separate color scales are shown because exponent and offset represent different quantities. In panels B–D, colors identify patients and marker shape identifies recording year.

Paired exponent and offset changes were then used to predict the frequency-resolved ON–OFF_mean_ contrast. The predicted contrast captured a substantial but variable fraction of the observed spectral change (median prediction *R*^2^ = 0.64 [IQR 0.50, 0.71]; Fig. 5A) and crossed zero at a median pivot frequency of 14.1 Hz [IQR 10.8–18.3 Hz].

Aperiodic correction changed the band-level interpretation (Fig. 5D). Raw theta decreased (median −0.83 dB), but adjusted theta was modestly positive (+0.17 dB), arguing against selective suppression. Raw low- and high-gamma increased in all sessions (medians +0.91 and +1.07 dB), yet residuals were much smaller: low gamma remained modestly positive (+0.12 dB; 7/10 sessions), whereas high gamma was inconsistent (−0.05 dB; 4/10 sessions). Exponent and offset changes were spatially widespread (Fig. 5E,F). Thus, the raw low-frequency decrease and much of the gamma increase reflected a broader aperiodic spectral-state shift, with a smaller residual most evident in low gamma.

## 4. Discussion

In this repeated-session proof-of-concept study, DBS ON produced a reproducible acute scalp-EEG state shift: raw gamma power increased in every session, lower-frequency differences depended on post-offset timing, and aperiodic exponent and offset decreased consistently. After aperiodic correction, theta suppression was not preserved, whereas a smaller low-gamma residual remained. The central result is therefore gamma enhancement embedded within a broader change in spectral shape, rather than an isolated band-limited effect.

These findings build on three lines of ANT-DBS literature: clinical studies establish therapeutic benefit but not acute scalp-EEG stimulation-state signatures [6–10]; invasive and network-level studies show modulation of hippocampal, thalamocortical, and broader epileptic networks, but rarely with repeated scalp EEG across yearly sessions [11, 34–37]; and prior scalp EEG studies report theta desynchronization, altered synchronization, and offset-locked responses, but not clearly the roles of OFF-window timing and aperiodic spectral structure [16–18]. We address the remaining gap by combining repeated within-patient recordings, time-resolved ON–OFF comparisons, STF mapping, and separation of aperiodic from residual spectral components.

The shortened stimulation paradigm was central to this design. The 1 s ON / 5 s OFF protocol was not intended to reproduce conventional 1 min ON / 5 min OFF therapeutic cycling [6–10]. Instead, it provided many repeated transitions within a short EEG session, enabling within-session ON–OFF comparisons and analysis of the first seconds after offset. The OFF-window analysis showed that this timing matters: low-frequency ON–OFF differences were strongest immediately after stimulation offset and weakened toward later OFF windows. The finding extends the offset-locked analysis of Mattila et al. [18] by showing that spectral effects evolve across the early post-offset interval.

The low-frequency findings are consistent with previous evidence that theta and other low-frequency dynamics are relevant to ANT-DBS network physiology. Scherer et al. [16] reported temporal theta-band desynchronization during effective ANT-DBS, and Aiello et al. [35] showed that theta-band anterior thalamus–cortex interactions contribute to stimulation-related network dynamics. Invasive studies have also linked ANT stimulation to hippocampal and seizure-network modulation [11, 34, 36]. The present results support the relevance of low-frequency dynamics, but refine their scalp-EEG interpretation: raw theta/alpha/beta decreases were not stable isolated band-limited effects, because they depended on post-offset timing and were strongly reduced after aperiodic correction. Thus, these low-frequency changes are better understood as part of a broader stimulation-state transition with early post-offset recovery, rather than as stable selective theta or alpha suppression.

The gamma-range finding fits the view of ANT-DBS as a network intervention and suggests that acute stimulation influences high-frequency activity detectable at the scalp. Gamma power increased during DBS ON in every session, with exploratory cross-year support for frontal low-gamma and central high-gamma patterns. A smaller low-gamma residual remained after aperiodic correction, indicating that the raw increase reflected both the broader spectral shift and a remaining band-limited component. This is biologically plausible because gamma activity has been linked to local circuit coordination and communication between neuronal populations [38, 39]. However, gamma rhythms, broadband gamma, and high-gamma activity can have different physiological origins [40], so the residual effect should be viewed as a candidate scalp-level feature rather than evidence for a single neural generator.

At first sight, our increase may seem different from reports of reduced beta/gamma power after chronic ANT-DBS and reduced 20–70 Hz activity during intraoperative stimulation [17, 35]. However, those studies examined different timescales, recording settings, and comparators, whereas we captured repeated acute ON–OFF transitions within the same recordings. Longitudinal intracranial ANT recordings provide a complementary timescale: responders developed progressive increases in β_1_/gamma and decreases in delta/theta activity over several years [12]. Such context-dependent effects are consistent with reports of hippocampal and temporal-lobe desynchronization and reduced large-scale epileptic-network coupling during stimulation [11, 16]. Further support comes from Liu et al. [41], who showed that ANT stimulation increased hippocampal gamma power and improved working-memory precision. Together, these observations support context-dependent ANT-DBS network effects rather than a direct contradiction between studies. Because our analysis was sensor-level, however, we cannot determine whether the residual low-gamma effect reflects cortical generators, thalamocortical interactions, hippocampal– anterior thalamic circuits, or mixed neural and non-neural sources.

The aperiodic results changed how the raw band-power effects were interpreted. Spectral parameterization showed that much of the apparent low-frequency decrease and gamma increase reflected a broader change in spectral shape: DBS ON consistently lowered the fitted aperiodic exponent and offset, and the aperiodic-predicted contrast reproduced a substantial portion of the frequency-dependent shape of the raw ON–OFF spectrum. Apparent low-frequency suppression may therefore partly reflect a flatter spectral background rather than selective theta or alpha suppression, illustrating the value of separating aperiodic background activity from periodic peaks [19]. A lower exponent may be consistent with a relative shift toward excitation [20, 42, 43], while recent neuromodulation studies show that aperiodic activity varies with DBS state and relates to seizure state or treatment response [21, 22, 44]. However, the slope– E/I relationship is context-dependent and not universally supported [45, 46]; sensor-level changes may also reflect firing dynamics, synaptic or membrane time constants, and spatial mixing. We therefore interpret the lower exponent conservatively as an acute spectral-state feature rather than a direct measure of cortical E/I balance.

Artifact-related explanations require explicit consideration because active DBS–EEG is susceptible to stimulation artifacts, line noise, and cranial muscle activity [23, 24, 47]. We therefore evaluated the gamma finding using three convergent robustness checks. First, continuous EEG was cleaned before cycle extraction, and ON and OFF windows were drawn from the same cleaned recording within each patient-year, ensuring matched recording and preprocessing histories. Second, the 140-Hz carrier and its integer harmonics were outside the analyzed 4–70 Hz range, while excluding 48–52 or 45–55 Hz retained 99.5% and 98.8% of significant positive STF nodes, respectively. Third, gamma support extended across a broad frequency range, showed spatial, temporal, and cross-year consistency, and a smaller low-gamma residual remained after aperiodic correction. This residual does not, by itself, establish a neural origin. Together, these checks argue against the stimulation carrier or narrowband 50-Hz contamination as sufficient explanations for the effect. They cannot exclude aliasing, spectral broadening, broadband muscle activity, or other stimulation-related contamination; source-level and dedicated artifact-focused validation therefore remain necessary.

The clinical relevance of this study is methodological and hypothesis-generating. Reproducible gamma enhancement may provide a noninvasive readout of acute stimulation-state engagement, while OFF-window dependence shows that reference timing can change ON–OFF interpretation. Aperiodic correction further shows that raw band power may overstate selective oscillatory effects. These points are relevant for future EEG-DBS studies comparing stimulation parameters, evaluating target-network engagement, and relating acute EEG responses to longer-term clinical outcomes. The present data do not establish clinical-response, cognitive, or outcome biomarkers, but define measurable acute EEG features that can be tested in those future contexts.

Several limitations define the next validation steps. The cohort was small (*n* = 5; four male and one female), limiting population-level inference and precluding sex-stratified analyses. No sham condition was included, and the within-cycle OFF windows were post-offset physiological comparators rather than an independent stimulation-free baseline. These constraints should be considered alongside the depth of this longitudinal clinical DBS–EEG dataset: 10 sessions recorded one year apart under unchanged stimulation programming, yielding 973 retained stimulation cycles. The repeated within-patient design supports cross-year consistency but does not replace independent validation. Repeated cycles may also have been temporally dependent, although the main PSD direction was preserved after cycle-block averaging. The analyses were sensor-level, so effects cannot be localized directly to ANT, hippocampal, cortical, or thalamocortical sources. The short 1 s windows limit low-frequency spectral resolution, and band-power and aperiodic estimates should be interpreted as short-window spectral-shape descriptions. Although stimulation contacts were anatomically localized relative to ANT, the small sample did not allow separation of the contributions of contact location, patient-specific anatomy, and seizure-network organization. Because all recordings used the same nominal bilateral voltage amplitude, amplitude-dependent effects could not be evaluated. Future studies should include head-model-based source localization, connectivity analysis, larger prospective cohorts, and clinical or cognitive follow-up to test behavioral or outcome relevance.

## 5. Conclusion

Across five ANT-DBS patients recorded at two time points, PSD, STF, and spectral-parameterization analyses identified reproducible acute scalp-EEG stimulation-state signatures. DBS ON was associated with increased raw gamma-range power and lower fitted aperiodic exponent and offset. STF showed cross-year frontal 34–44 Hz and central 56–70 Hz support. After aperiodic correction, a modest low-gamma residual remained, whereas low-frequency suppression was not preserved and depended on OFF-window timing. Overall, these findings suggest that gamma-range enhancement occurred within a broader stimulation-related spectral-state shift and define candidate acute EEG signatures for future source-level, connectivity-based, and prospective validation studies.

## CRediT authorship contribution statement

Elaheh Sabbaghi: Conceptualization, Methodology, Software, Formal analysis, Visualization, Writing – original draft.

Herkko Mattila: Data curation, Investigation, Resources, Methodology, Writing – review & editing.

Eetu Siitama: Visualization, Writing – review & editing.

Jari Hyttinen: Supervision, Methodology, Writing – review & editing.

Jukka Peltola: Resources, Supervision, Writing – review & editing.

Narayan Puthanmadam Subramaniyam: Supervision, Conceptualization, Writing – review & editing.

All authors read and approved the final manuscript.

## Ethics approval and consent to participate

The study was conducted in accordance with the Declaration of Helsinki and was approved by the Ethics Committee of the Pirkanmaa Hospital District (decision R17132; approved 19 September 2017). Written informed consent was obtained from all participants.

## Consent for publication

Not applicable.

## Availability of data and materials

De-identified EEG data are not publicly available due to patient privacy and institutional restrictions. Research data are conditionally available from the authors under a formal data sharing agreement and appropriate approvals.

## Code availability

The analysis code supporting the findings of this study will be made publicly available when the manuscript enters external peer review at https://github.com/elahehsabbaghi/ant-dbs-eeg-acute-signatures.

## Supplementary material

Supplementary material associated with this article is available in the online version.

## Acknowledgements

We thank the clinical neurophysiology and epilepsy-DBS teams at Tampere University Hospital for support with EEG acquisition and clinical coordination. We also thank members of the FAME/DREAM project and doctoral school community for helpful discussions and support.

## Declaration of Generative AI and AI-assisted technologies in the manuscript preparation process

During the preparation of this work, the authors used ChatGPT (OpenAI) to support language editing and manuscript organization. After using this tool, the authors reviewed and edited the content as needed and take full responsibility for the content of the published article.

