## Supplementary Material for "Acute scalp EEG signatures of anterior thalamic DBS in epilepsy: reproducible gamma power enhancement within an aperiodic spectral-state shift"

**For:**

### Scope of this supplement

This supplement provides implementation details and selected supporting cohort/stimulation information that were abbreviated in the main text. Notation follows the main text: $t_0$ denotes stimulation offset, and ON/OFF windowing follows the nominal 1 s ON / 5 s OFF protocol.

### Supplementary cohort and stimulation details

DBS programming parameters were identical in the 2018 and 2019 recordings and are summarized in Table S1. A descriptive group-level visualization of reconstructed DBS leads and contacts in MNI space together with the DISTAL atlas representation of the bilateral ANT is shown in Fig. S1.

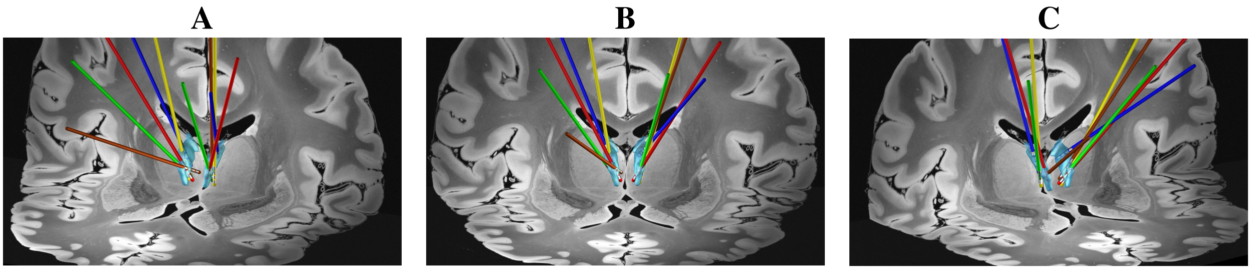


Figure S1: **Descriptive group-level visualization of DBS leads and contacts.** DBS lead trajectories and contact locations from five patients are shown in normalized MNI space relative to the DISTAL atlas representation of the bilateral ANT. Panels A–C show the same group-level visualization from different viewing angles. Colors indicate individual patients: P01, red; P02, blue; P03, green; P04, yellow; and P05, brown. The figure is provided for anatomical context only, and no quantitative spatial analysis was performed.

### Supplementary data yield

Quality-control yield, including candidate cycles, rejected cycles, retained cycles, and channel availability, is summarized in Table S2. These counts reflect the preprocessing, channel-quality flagging, and window-validity criteria described in the Supplementary Methods below.

### Supplementary Methods

#### EEG acquisition and stimulation details

EEG acquisition followed the ANT-DBS framework described by Mattila et al. [1]. EEG was recorded using a 64-channel active-electrode cap (actiCAP slim or actiCAP snap, Brain Products GmbH, Gilching, Germany) arranged according to the international 10–10 system. The recording protocol targeted electrode impedances below 5 kΩ, FCz served as the recording reference, and Fpz served as ground. Two bipolar electrooculography channels, for horizontal and vertical eye movements, and a two-lead electrocardiography channel were recorded concurrently. All electrodes were connected to a NeurOne Tesla amplifier (Bittium Biosignals, Oulu, Finland). Signals were sampled at 20 kHz; the

**Table S1**
**DBS stimulation settings used during the analyzed 2018 and 2019 DBS–EEG recordings.**

| Patient | Device | Configuration | Stimulated contacts L/R | Voltage amplitude L/R (V) |
|---|---|---|---|---|
| P01 | Medtronic Activa PC | Bilateral monopolar | -3 / -11 | 5.0 / 5.0 |
| P02 | Medtronic Activa PC | Bilateral monopolar | -3 / -11 | 5.0 / 5.0 |
| P03 | Medtronic Activa PC | Bilateral monopolar | -3 / -11 | 5.0 / 5.0 |
| P04 | Medtronic Activa PC | Bilateral monopolar | -3 / -11 | 5.0 / 5.0 |
| P05 | Medtronic Activa PC | Bilateral monopolar | -2 / -11 | 5.0 / 5.0 |

*Note.* All patients had bilateral Medtronic 3389 DBS leads connected to Medtronic Activa PC implantable pulse generators. DBS programming settings were unchanged between the 2018 and 2019 recordings. All patients received voltage-controlled bilateral monopolar stimulation at 5.0 V per side, 140 Hz, and 90 $\mu$s pulse width. The experimental protocol used nominal 1 s ON / 5 s OFF cycling for approximately 10 min. L/R indicates left/right stimulation settings.

**Table S2**
**Quality-control yield by patient-year session.** Candidate cycles denote extracted stimulation cycles available for analysis. Retained cycles entered the main ON–OFF$_{mean}$ analyses and required a valid 1 s ON window and an OFF$_{mean}$ estimate based on at least 3 of 5 valid OFF windows. Rejected cycles failed these window-validity criteria. Five-cycle blocks denote non-overlapping temporal blocks used in the supplementary cycle-block sensitivity analysis; the final block was retained if it contained at least three cycles. Excluded and valid channel counts are summarized across retained cycles as median [range]; a single number indicates no variation across retained cycles.

| Session | Candidate cycles | Rejected cycles | Retained cycles | 5-cycle blocks | Excluded channels | Valid channels |
|---|---|---|---|---|---|---|
| P01 2018 | 93 | 0 | 93 | 19 | 0 | 64 |
| P02 2018 | 95 | 0 | 95 | 19 | 0 | 64 |
| P03 2018 | 92 | 0 | 92 | 18 | 0 | 64 |
| P04 2018 | 97 | 0 | 97 | 19 | 0 | 64 |
| P05 2018 | 90 | 0 | 90 | 18 | 0 | 64 |
| P01 2019 | 99 | 1 | 98 | 20 | 0 [0–1] | 64 [63–64] |
| P02 2019 | 103 | 1 | 102 | 20 | 1 [1–2] | 63 [62–63] |
| P03 2019 | 100 | 1 | 99 | 20 | 0 | 64 |
| P04 2019 | 103 | 2 | 101 | 20 | 0 [0–1] | 64 [63–64] |
| P05 2019 | 106 | 0 | 106 | 21 | 0 | 64 |
| Total | 978 | 5 | 973 | 194 | – | – |

*Note.* Cycle rejection was minimal and occurred only because a small number of extracted epochs were shorter than the required analysis span. Channel counts reflect analysis-level channel availability after preprocessing and channel-quality flagging. Five-cycle block counts refer to the supplementary temporal-dependence sensitivity analysis and were computed from retained cycles only.

amplifier input frequency range was DC–3500 Hz and the channel dynamic range was ±430 mV. Brainstorm software (version 20-May-2019) [2] was used for the original visual inspection and signal-preprocessing workflow.

All patients had bilateral Medtronic 3389 DBS leads connected to Medtronic Activa PC implantable pulse generators. The analyzed EEG condition used voltage-controlled bilateral monopolar ANT stimulation at 5.0 V per side, 140 Hz, and a pulse width of 90 $\mu$s. Left- and right-sided pulses were delivered alternately rather than simultaneously, so each side was stimulated at 140 Hz, with a combined alternating bilateral pulse stream of 280 pulses/s. Stimulation followed a nominal 1 s ON / 5 s OFF protocol for approximately 10 min. DBS programming settings were unchanged between the 2018 and 2019 recordings and are summarized in Table S1.

### DBS lead localization and visualization

DBS electrodes were localized using the default pipeline implemented in Lead-DBS v3 [3–5]. MATLAB R2023b was used for the Lead-DBS analysis pipeline. Postoperative CT images were linearly coregistered to preoperative T1-weighted MRI images using Advanced Normalization Tools [6]. The preoperative MRI was then nonlinearly normalized to the ICBM 2009b Nonlinear Asymmetric MNI template using the multi-stage SyN-based procedure with subcortical refinement implemented in the default DBS pipeline in Lead-DBS [7, 8].

To reduce the effect of intraoperative brain shift, a subcortical-restricted affine refinement between preoperative and postoperative images was applied using the Lead-DBS brain-shift correction module [9]. All preprocessing steps were visually inspected and manually adjusted when necessary. If clear mismatch was observed, normalization was manually refined using WarpDrive [10].

After preprocessing, electrodes were pre-localized using the PaCER algorithm [11]. In cases where automatic reconstruction was insufficient, TRAC/CORE and/or manual refinement of trajectories and contact positions in the Lead-DBS interface was used. For anatomical context, reconstructed electrodes and contacts were visualized in MNI space together with the bilateral ANT labels from the DISTAL subcortical atlas [12]. Group-level visualization was performed using the Lead Group module in Lead-DBS [4]. Electrode reconstructions were included to document the anatomical distribution of the implanted leads and to show that their trajectories were broadly consistent with the intended ANT region. The visualization was descriptive; no quantitative assessment of ANT coverage, contact-to-target distance, stimulation-volume overlap, or associations between electrode location and EEG responses or clinical outcomes was performed.

### EEG preprocessing and offset-locked windowing

Raw EEG preprocessing followed the pipeline of Mattila et al. [1] and was performed in Brainstorm (version 20-May-2019). Continuous recordings were visually screened before artifact correction. For each recording session, 64 independent components were estimated from the continuous data using the InfoMax algorithm after resampling the ICA input to 2 kHz. Components were selected for removal by visual evaluation of their time series and scalp topographies and by inspection of the EEG distortion produced by their removal. This procedure was used to attenuate ocular, small-amplitude muscle, and stimulation-related artifacts. The cleaned continuous EEG was retained at 2 kHz for subsequent cycle extraction.

ICA component-exclusion lists and channel-quality flags were defined separately for each recording session before extracting DBS ON and OFF windows. ON and OFF windows were therefore obtained from the same session-level cleaned continuous recording, rather than being cleaned separately by stimulation state.

The source datasets had slightly different prior filtering histories across recording years: the 2018 recordings had been band-pass filtered over 0.5–100 Hz, whereas the 2019 recordings had been filtered over 0.5–80 Hz with an additional 50 Hz notch. For the present analyses, residual 50 Hz line noise was attenuated uniformly across all patient-year sessions using the MNE `spectrum_fit` notch implementation. Each 1-s ON or OFF analysis window was subsequently rereferenced to the common average of the channels marked valid for that cycle before spectral estimation. All reported spectral analyses were restricted to 4–70 Hz, below the 2019 low-pass cutoff. Although the prior filtering difference can remain visible in absolute PSD plots, within-session ON–OFF comparisons shared the same prior filtering history.

Stimulation offset ($t_0 = 0$), defined as the last pulse of each nominal 1 s burst, was used as the temporal anchor. Each extracted epoch contained 14,001 samples at 2,000 Hz and spanned $-1.50$ to 5.50 s. The ON window covered $-0.98$ to 0.02 s. After a 120 ms guard interval, five consecutive 1 s OFF windows were analyzed: $\text{OFF}_1 = 0.12$–1.12 s, $\text{OFF}_2 = 1.12$–2.12 s, $\text{OFF}_3 = 2.12$–3.12 s, $\text{OFF}_4 = 3.12$–4.12 s, and $\text{OFF}_5 = 4.12$–5.12 s. Event markers placed the next stimulation onset at approximately 5.132 s, 12 ms after $\text{OFF}_5$ ended. Thus, none of the analyzed OFF windows overlapped the following stimulation burst. The 7 s value refers to the extracted epoch length, not to the interval between stimulation bursts.

$\text{OFF}_{\text{mean}}$ was calculated when at least 3 of the 5 OFF windows were valid. Retained cycles required a valid ON window and at least 3 valid OFF windows. Overall, 973 of 978 candidate cycles were retained; five were excluded because their extracted epochs were shorter than the required analysis span.

Because stimulation-related artifacts can appear at the stimulation frequency, its harmonics, or at lower frequencies through aliasing or spectral broadening, the main findings were interpreted together with aperiodic correction, cross-year replication, STF spatial–temporal patterns, and explicit artifact-related limitations rather than from raw band power alone [13, 14].

### PSD estimation

Power spectra were estimated separately for each retained channel and 1 s analysis window using multitaper discrete prolate spheroidal sequence estimation in MNE-Python. Spectra were computed over 4–70 Hz using MNE-Python's `psd_array_multitaper` function with adaptive weighting disabled, full normalization, and the default bandwidth setting. With the 1 s windows used here, the default bandwidth corresponds to approximately 8 Hz spectral smoothing, equivalent to a time-half-bandwidth product of approximately 4. No explicit zero-padding was applied; therefore, the native frequency grid had approximately 1 Hz spacing. Multitaper spectra were averaged in linear power units before conversion to decibels.

For PSD summaries, channel-wise spectra were converted to dB and averaged across retained channels within each window. ON–OFF contrasts were then computed as paired within-cycle differences in dB. For sensor-level cluster-permutation testing, the channel-by-frequency dB spectra were retained before forming paired ON–OFF contrasts. Band summaries used non-overlapping computational boundaries: theta, $4 \leq f < 8$ Hz; alpha, $8 \leq f < 12$ Hz; beta, $12 \leq f < 30$ Hz; and gamma, $30 \leq f \leq 70$ Hz.

### Cycle-block sensitivity analysis

To assess whether the primary PSD findings were sensitive to repeated-cycle dependence, we performed a supplementary cycle-block analysis. Within each patient-year session, consecutive retained cycles were averaged into non-overlapping 5-cycle temporal blocks before recomputing the $\mathrm{ON{-}OFF_{mean}}$ band-power contrast. The final block was retained if it contained at least three cycles. Session-level effects were then summarized from block-level mean contrasts. This analysis was used as a robustness check for temporal dependence between adjacent stimulation cycles and was not treated as a replacement for the primary within-session cluster-permutation analysis.

### Spatiotemporal–frequency (STF) decomposition

Spatiotemporal–frequency (STF) decomposition described how DBS-related spectral changes evolved across sensors, frequencies, and time within each 1 s window. Time–frequency representations were computed for each channel and cycle using complex Morlet wavelets from 4 to 70 Hz in 1 Hz steps [15]. The number of wavelet cycles increased with frequency,

$$n_{\text{cycles}} = \max(1,\ 0.35f)\,,$$

and was reduced iteratively when needed so that the effective kernel fit within the 1 s window.

Power was converted to dB and summarized on a common grid of 20 frames from 0.00 to 0.95 s in 50 ms steps. In the ON segment, time 0 corresponds to the start of the extracted window (−0.98 s relative to stimulation offset); in each $\mathrm{OFF}_k$ segment, time 0 corresponds to the start of that OFF window. Morlet outputs were mapped to the common grid by interpolation. This grid standardizes visualization and should not be interpreted as additional temporal precision.

*Line-noise profiles.* Residual 50-Hz line noise was attenuated before STF decomposition using multitaper spectral fitting. The primary `notch_only` analysis retained the complete 4–70 Hz inferential domain. Two additional sensitivity profiles repeated the full cluster analysis after excluding 48–52 Hz or 45–55 Hz before cluster formation. Excluded frequency rows remained in the saved frequency axis but were removed from the inferential node mask and disconnected from the adjacency graph, preventing clusters from bridging the gap. The three profiles used identical notch-cleaned TFR data, cycle ordering, and random seed, but each profile had its own maximum-cluster permutation null distribution.

*Regions and frequency summaries.* Macro-region assignment was applied to the first 64 scalp EEG channels: frontal = Fp*, AF*, F* (excluding FC* and FT*); central = FC*, C*, CP*; temporal = T*, FT*, TP*; parietal = P*, PO*; and occipital = O* and Iz. These regions were used only for descriptive summaries and did not define the sensor–frequency–time cluster tests. Conventional bands were theta, $4 \leq f < 8$ Hz; alpha, $8 \leq f < 12$ Hz; and beta, $12 \leq f < 30$ Hz. Line-noise-robust gamma summaries and candidate searches used 30–44 and 56–70 Hz. The primary atlas and regional maps retained the complete notch-cleaned 4–70 Hz spectrum; 48–52 Hz was marked lightly for display only.

*STF visualization summaries.* For the patient-year atlas, within-session $t$-statistic maps were averaged across sensors and $\mathrm{ON{-}OFF_1}$ through $\mathrm{ON{-}OFF_5}$. The cohort map was the median across the 10 patient-year maps, and sign consistency was the number of positive minus negative patient-years at each bin. Atlas and cohort maps were smoothed for display only ($\sigma = 0.75$ frequency bins and 0.45 time bins); quantitative summaries used unsmoothed values. The descriptive safe-gamma value was the mean $t$ statistic over sensors, OFF references, time bins, and the union of 30–44 and 56–70 Hz.

Cross-year signed-support maps show the fraction of patients with replicated ON>OFF support minus the fraction with replicated ON<OFF support. Open circles mark sensors with $|\text{support}| \geq 3/5$. For OFF-reference sensitivity,

support at each bin was the proportion of patient-year sessions with at least one significant sensor of the specified sign. Values were averaged over 30–44 and 56–70 Hz for gamma ON>OFF, or over 4–15 Hz for low-frequency ON<OFF.

*Candidate ROI discovery.* Candidate ROIs were derived from the primary `notch_only` profile. For each patient and year, stable sign-specific masks required the same sign in at least 3 of 5 ON–OFF$_k$ contrasts and no opposite-sign membership. A regional bin was counted as supported only when at least

$$\max\left(2, \lceil 0.10 N_{\text{region}} \rceil\right)$$

sensors in that region showed stability in both years. Patient maps were then summed, and candidate bins required cross-year prevalence in at least 3/5 patients. Connected components were identified separately within 30–44 and 56–70 Hz using four-neighbor frequency–time adjacency. Components had to span at least three frequency bins and five 50-ms time bins. Within each region, band, and direction, components were ranked by peak prevalence and component area. The complete table was ordered by peak prevalence, ICC, and median $|t|$, and the main figure shows the highest-ranked positive candidate in each safe-gamma interval.

For each fixed ROI, ICC(3,1) summarized consistency between 2018 and 2019 regional support-prevalence values. Median $|t|$ was calculated over all regional sensors, ROI bins, years, and OFF references for each patient and then summarized across patients. Candidate generation and stability estimation used the same dataset; these values are therefore post-selection descriptive summaries.

### STF line-noise sensitivity results

The guarded profiles closely reproduced the primary notch-only pattern. In the common 30–44 and 56–70 Hz domain, the 48–52 and 45–55 Hz profiles retained 99.54% and 98.77%, respectively, of notch-only significant positive nodes. Safe-gamma support was present in 50/50 patient-year/OFF comparisons in the notch-only and 48–52 Hz profiles and in 49/50 comparisons under the conservative 45–55 Hz guard. Cross-year replication was similarly stable across profiles.

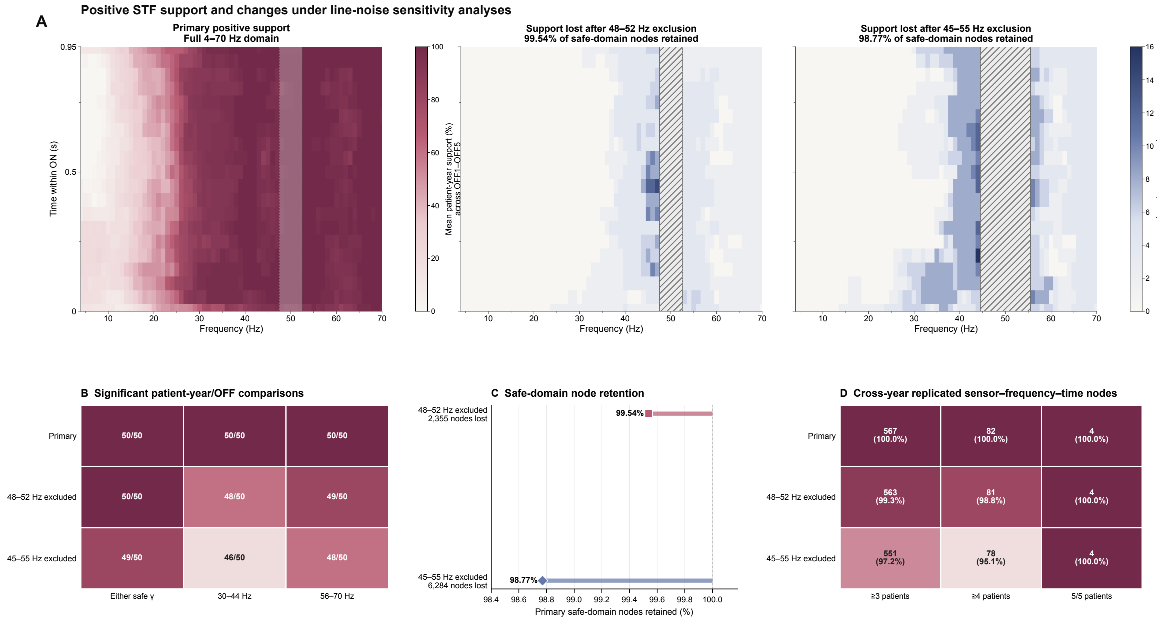


Figure S2: **Sensitivity of positive STF support to line-noise handling.** (A) The primary map shows mean positive patient-year support across ON–$\mathrm{OFF}_1$ through ON–$\mathrm{OFF}_5$. The two sensitivity maps show the reduction in mean support relative to the primary analysis after excluding 48–52 or 45–55 Hz before cluster formation. Hatched areas mark the excluded frequency intervals. (B) Number of patient-year/OFF comparisons with significant positive support in either line-noise-safe gamma interval, 30–44 Hz, or 56–70 Hz. (C) Percentage of primary positive nodes retained in the common safe-gamma domain, together with the number of nodes lost. (D) Number of sensor–frequency–time nodes showing cross-year replication in at least three patients, at least four patients, or all five patients. Percentages are relative to the primary analysis within each replication threshold.

### Aperiodic parameterization and aperiodic-adjusted metrics

To separate the fitted aperiodic component from residual spectral structure, each cycle- and channel-specific spectrum was parameterized with `specparam` using a fixed aperiodic model [16]:

$$\log_{10} P(f) = b - \chi \log_{10}(f),$$

where $b$ denotes the offset and $\chi$ the aperiodic exponent. Throughout the manuscript, changes are reported as $\Delta\chi = \chi_{\mathrm{ON}} - \chi_{\mathrm{OFF_{mean}}}$ and $\Delta b = b_{\mathrm{ON}} - b_{\mathrm{OFF_{mean}}}$. The equivalent log–log slope is $-\chi$.

Models were fitted over 4–70 Hz after linear interpolation in $\log_{10}$-power space across 49–51 Hz. A maximum of six periodic peaks was allowed, with peak-width limits of 2–12 Hz, a minimum peak height of 0.05, and a peak threshold of 2.0. These settings were chosen to support stable estimation of the aperiodic component rather than exhaustive characterization of periodic peaks.

Channel-by-cycle estimates were retained when the coefficient of determination of the fitted aperiodic line over 4–70 Hz was at least 0.20. The full `specparam` model $R^2$ was retained for quality-control reporting but was not used as the primary filtering criterion. Fits failing the aperiodic-line $R^2$ criterion were set to missing on a channel-by-cycle basis. Sensor-level analyses required at least 10 valid paired cycle estimates per channel. No additional cycle-level exclusion based on a fixed post-fit channel-count threshold was applied.

For each cycle and channel, $\mathrm{OFF_{mean}}$ was calculated only when at least three of the five OFF estimates remained valid after model fitting and fit-quality filtering. This reapplied the three-of-five OFF-window requirement after channel-level model quality control.

Aperiodic-adjusted spectra were calculated as

$$\log_{10} P_{\mathrm{adj}}(f) = \log_{10} P_{\mathrm{interp}}(f) - \left[b - \chi \log_{10}(f)\right],$$

where $P_{\text{interp}}(f)$ denotes the spectrum after interpolation of the 49–51 Hz region.

Raw band metrics were calculated from the uninterpolated spectra, whereas adjusted metrics were calculated from $P_{\text{adj}}(f)$. Non-overlapping computational boundaries were used for theta ($4 \leq f < 8$ Hz), low-gamma ($30 \leq f < 55$ Hz), high-gamma ($55 \leq f \leq 70$ Hz), and broad gamma ($30 \leq f \leq 70$ Hz). Frequencies from 49 to 51 Hz were excluded from gamma-range averaging. $\text{Log}_{10}$-power differences were multiplied by 10 for reporting in decibels. These quantities describe residual band-limited spectral effects after removal of the fitted aperiodic component.

For the frequency-resolved prediction analysis shown in the main manuscript, the aperiodic-predicted contrast within each patient-year was calculated from the paired offset and exponent differences:

$$\Delta\widehat{\log_{10} P}(f) = \Delta b - \Delta\chi \log_{10}(f).$$

Observed and predicted contrasts were calculated from the same fit-quality-qualified channel–cycle observations. Values were averaged across matched valid channels within each cycle and then across cycles within each patient-year. The residual curve was calculated as the observed contrast minus the predicted contrast. Patient-year curves were aligned to a common 1-Hz frequency grid before calculating the cohort median and interquartile range. The aperiodic pivot frequency was calculated as

$$f_{\text{pivot}} = 10^{\Delta b/\Delta\chi}$$

when the resulting value lay within the fitted frequency range.

Prediction $R^2$ quantified agreement across frequency bins between the observed and aperiodic-predicted ON–$\text{OFF}_{\text{mean}}$ curves within each patient-year, excluding 49–51 Hz. This measure was distinct from both the aperiodic-line and full-model fit-quality $R^2$ values. Sensitivity analyses repeated the summaries using full-model $R^2 \geq 0.20$, both aperiodic-line and full-model criteria, or all successful finite fits. These alternatives produced the same overall interpretation as the primary aperiodic-line $R^2 \geq 0.20$ analysis.

### Sensor adjacency

For cluster-based permutation testing, sensor adjacency was defined from the recorded electrode coordinates by linking EEG sensor pairs whose Euclidean distance was ≤5 cm (first 64 EEG channels). This threshold produced a local neighborhood graph on the digitized montage, preserving nearby scalp contiguity while limiting long-range bridges during cluster formation. For visualization only, topographies were shown using a standard cap layout (Brain Products RNP-BA-128); this had no effect on the statistical adjacency structure or the results.

### Statistical analysis and multiplicity control

Given the small cohort and the large number of repeated cycles per recording, statistical testing was performed separately within each patient-year, with cycles treated as paired within-session observations. Across patient-years, we report consistency rather than pooled group-level inference. Because adjacent cycles may be temporally correlated, sign-flip testing assumes only approximate exchangeability of paired differences under the null. Results are therefore interpreted through within-session evidence and cross-year replication, without treating the number of cycles as a population-level sample size.

All statistical tests were based on within-session paired contrasts between ON and the corresponding OFF segment(s). We used nonparametric sign-flip permutation testing with cluster-based correction for multiple comparisons in channel×frequency (PSD), channel×frequency×time (STF), or channel space (aperiodic and derived metrics), following established methods [17–19]. Permutation distributions were generated by Monte Carlo sign-flip resampling with $N_{\text{perm}} = 5000$ permutations per patient-year and analysis family (PSD, STF, aperiodic/derived metrics, and OFF-progression slopes), providing a practical balance between stable cluster-level $p$-value estimation and computational tractability. Cluster-level $p$-values were computed as

$$p = \frac{1 + \sum_{r=1}^{N_{\text{perm}}} \mathbb{I}\left(M_r \geq M_{\text{obs}}\right)}{N_{\text{perm}} + 1},$$

where $M_r$ is the maximum absolute cluster mass obtained in permutation $r$, and $M_{\text{obs}}$ is the absolute mass of the observed cluster. Full enumeration of all $2^n$ sign-flip patterns was not used because the number of paired cycles per patient-year made exhaustive evaluation impractical.

*Paired contrasts and inclusion minima.* For each patient-year, per-cycle paired differences ($\text{ON}-\text{OFF}_k$) were formed for each feature and tested against zero using two-sided one-sample tests across cycles. Testing was performed only when at least 10 paired cycles were available for PSD and aperiodic-derived metrics, and at least 8 for STF. These minima were used as pragmatic stability requirements for within-session permutation testing; the lower STF threshold reflects additional loss of usable paired observations near window boundaries due to wavelet-support constraints. For $\text{OFF}_{\text{mean}}$-based summaries, $\text{OFF}_{\text{mean}}$ was computed within a cycle only when at least 3 of 5 OFF windows were valid, to avoid unstable baselines caused by sparse OFF-slice availability. For aperiodic metrics, this three-of-five criterion was reapplied separately for each channel after model fitting and fit-quality filtering.

*Scalar aperiodic summaries.* For each patient-year, the mean cycle-level $\text{ON–OFF}_{\text{mean}}$ difference and paired effect size $d_z$ were calculated for exponent, offset, and raw or aperiodic-adjusted band metrics. Across the 10 patient-year sessions, these quantities were summarized descriptively using the median [IQR], the number of sessions with $\Delta > 0$, cross-year replicated sign, and the median $d_z$.

*Cluster formation and statistics.* Clusters were defined using the appropriate sensor, frequency, and time adjacency. At every tested node, the $t$ statistic, degrees of freedom, and two-sided uncorrected $p$ value were calculated using the valid paired cycles available at that node. Cluster-forming nodes required $p < 0.05$ and at least 10 valid pairs for PSD and aperiodic-derived metrics or eight for STF. For primary STF inference, all 4–70 Hz nodes remained eligible after notch attenuation. In the two sensitivity profiles, 48–52 or 45–55 Hz nodes were excluded before cluster formation while remaining in the saved frequency axis; adjacency edges touching the excluded nodes were removed. In the aperiodic analyses, this channel-wise cluster framework was applied to sensor-space maps of derived features such as exponent, offset, aperiodic-adjusted band metrics, and related OFF-progression summaries. Cluster mass was defined as the sum of $|t|$ values within each same-sign cluster, and cluster significance was based on |mass|.

*FWER across $OFF_1$–$OFF_5$ (max-statistic).* The primary $\text{ON–OFF}_{\text{mean}}$ PSD and aperiodic contrasts used single-contrast maximum-cluster null distributions. For $\text{ON–OFF}_1$ through $\text{ON–OFF}_5$, the maximum absolute cluster mass across all five OFF references was retained in each permutation and used to evaluate every $\text{OFF}_k$ cluster, controlling family-wise error across OFF references at $\alpha = 0.05$.

*OFF progression slopes.* To summarize systematic post-offset evolution, we computed per-cycle slopes across $\text{OFF}_1$–$\text{OFF}_5$ for selected features by regressing feature values against the mid-time of each OFF window. Given $\text{OFF}_k = [0.12 + (k-1),\ 1.12 + (k-1)]$ s, the corresponding mid-times are 0.62, 1.62, 2.62, 3.62, and 4.62 s. Slopes were computed only when at least four OFF windows were valid for the cycle and channel, allowing occasional missing slices while still requiring most OFF windows for a stable linear trend estimate. Within each patient-year, slope maps were evaluated using one-sample sign-flip cluster permutation testing in sensor space with $p$-based cluster formation ($\alpha = 0.05$); across patient-years, we summarize directional consistency. For the band-level PSD summaries reported in the main text, OFF-progression slopes were computed separately at the patient-year level by regressing $\text{ON–OFF}_k$ band differences against OFF-slice index ($k = 1, \ldots, 5$); these slopes are therefore reported in dB/slice.
